# Incremental Accumulation of Linguistic Context in Artificial and Biological Neural Networks

**DOI:** 10.1101/2024.01.15.575798

**Authors:** Refael Tikochinski, Ariel Goldstein, Yoav Meiri, Uri Hasson, Roi Reichart

**Affiliations:** The Faculty of Data and Decisions Sciences, Technion - Israel Institute of Technology, Haifa, Israel.; Department of Cognitive and Brain Sciences, The Hebrew University of Jerusalem, Jerusalem, Israel.; Department of Psychology, Princeton University, Princeton, NJ, USA

## Abstract

Accumulated evidence suggests that Large Language Models (LLMs) are beneficial in predicting neural signals related to narrative processing. The way LLMs integrate context over large timescales, however, is fundamentally different from the way the brain does it. In this study, we show that unlike LLMs that apply parallel processing of large contextual windows, the incoming context to the brain is limited to short windows of a few tens of words. We hypothesize that whereas lower<u>-</u>level brain areas process short contextual windows, higher-order areas in the default-mode network (DMN) engage in an online incremental mechanism where the incoming short context is summarized and integrated with information accumulated across long timescales. Consequently, we introduce a novel LLM that instead of processing the entire context at once, it incrementally generates a concise summary of previous information. As predicted, we found that neural activities at the DMN were better predicted by the incremental model, and conversely, lower-level areas were better predicted with short-context-window LLM.

## Introduction

Recent works in cognitive neuroscience have demonstrated the power of computational Large Language Models (LLMs) in predicting language-evoked neural signals among humans.^1–10^ LLMs have revolutionized the natural language processing (NLP) field, demonstrating human-and super-human-level performance in many language tasks.^11–13^ These models are capable of producing a rich language representation, manifested via multidimensional numerical vectors, also known as word embedding representations. The representations are also contextualized as the embedding representation of a word may be changed according to the context in which it appears, namely the preceding words in the input text. A growing number of studies show that these contextual representations can be linearly mapped to neural signals (e.g., fMRI, EEG, ECoG) recorded from human participants listening to spoken narratives – an analysis commonly referred to as neural encoding.^4,7,8,10^ In this method, a contextualized word embedding vector is extracted for each word in the narrative by providing the LLM with that word, along with a *context window* of the *N* preceding words (Figure 1c). The extracted vectors then serve as the input to a linear regression model that predicts the neural signal evoked by the corresponding words. The success of the neural encoding method, together with the human-like cognitive mechanism that leverages contextual information in the LLM, suggests that LLMs could serve as a plausible cognitive model for how language is computed in the human mind.^10,14,15^

**Figure 1.**
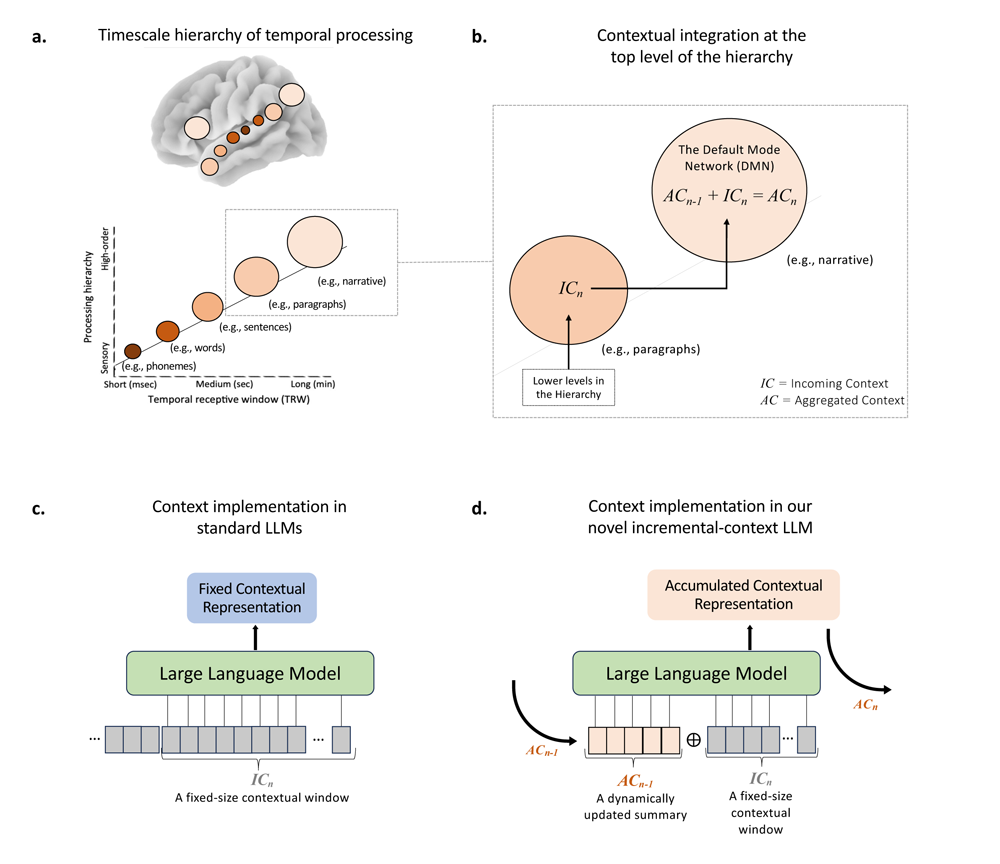
**a**. The neuroanatomical hierarchical organization according to multiple timescales of processing. Adapted from^18^. **b.** Our proposed neural mechanism of integrating long-term contextual information at the top level of the timescale hierarchy. **c.** The baseline implementation of contextual integration via Large Language Models (LLMs). The model is exposed to the entire incoming context window and processes it in a parallel manner. **d.** Our proposed alternative model of contextual integration via LLM. Instead of processing the entire context window at once, the incremental-context LLM is applied sequentially along the story. The LLM accumulates long-term contextual information by generating a concise summary of the past, and at each step, integrating this summary with the incoming context window and updating the summary to be used in the next step (see more details in Figure 3 and S3).

One of the drawbacks in considering LLMs as cognitive models, however, is manifested in the way these models process natural texts, such as stories or narratives, which unfold over long timescales. In contrast to the human brain, LLM digests large body of text, comprising of thousands of words in parallel, using a fixed-size contextual window. Thanks to the underlined attention mechanism,^16^ the LLM can learn the contextual dependencies across all words in parallel. In contrast, the human brain processes the incoming linguistic input serially, word by word, as speech and text unfold over time. Furthermore, humans don’t remember or have direct access to every single word they processes but rather seem to operate an online mechanism for accumulating information and integrating it into a broader contextual memory that is changed and updated as the story unfolds.^17,18^ In this study, we aim to provide an alternative model for how the brain, as opposed to current LLMs, integrates linguistic information over short-term and long-term contexts.

A series of studies showed that the brain gradually integrates temporal information across cortical areas in a topographic hierarchical manner. In such a topography, the temporal receptive windows (TRW) gradually increase along the cortical processing hierarchy, with early sensory areas integrating speech-related information (e.g., phonemes) into words over short periods of time (tens to hundreds of milliseconds). Adjacent cortical regions then integrate word-level information into sentences over several seconds and transfer the information into adjacent areas, which integrate the sentences into paragraphs. Finally, areas along the default mode network (DMN), located at the top of the temporal integration hierarchy, can integrate the paragraphs into a coherent narrative by integrating information accumulated over hundreds of seconds as the story unfolds, with relevant past information stored in long-term memories (Figure 1a).^18–20^ Such temporal processing hierarchical topography provides an alternative processing scheme for integrating short-term and long-term linguistic information over time within the DMN (Figure 1b).

We hypothesize that unlike LLMs which process large contextual windows of thousands of words, DMN networks can receive information about the incoming context (IC) through a small window of just tens of words (Fig. 1B). To test this hypothesis, a group of participants listened to several spoken stories while undergoing fMRI scans (a total of 297 scans recorded from 219 individuals). We then designed and implemented several encoding models to predict their neural responses using contextual embeddings extracted from an LLM after parametrically adjusting the contextual window size of the LLM from just a few words to a thousand words. We demonstrate empirically that the fit between the LLM and the brain decreases as the size of the LLM’s context window increases beyond tens of words, and that the maximal fit is obtained when the context window size is ∼32 tokens in length. This result supports our prediction that the incoming contextual information to the brain integrates information over a few sentences.

Next, we hypothesize that incoming contextual information at time *n* (IC_n_) is integrated with the aggregated contextual information (AC_n-1_) already accumulated in the DMN (Figure 1B). At the beginning of a story, where no contextual information has been accumulated yet, the accumulated context matches the incoming context (AC_1_=IC_1_). As the story unfolds, the accumulated contextual information is the sum of the incoming and aggregated contextual information. To test this prediction, we suggested an alternative LLM-based incremental-context model that fuses the incoming short-term context (IC_n_) with the aggregated contextual (AC_n-1_). The aggregated prior context is operationalized by asking the LLM to generate a concise summary of the incoming contextual information—a summary that is incrementally changed and updated as the model progresses through the narrative (see Figures 1d, 3b, S3, and Method). Adding a summary of the aggregated contextual information to the incoming information greatly improved our ability to predict the neural responses in the brain while processing all narratives, and this improvement was mainly evident in the higher-order areas among the DMN. Combined, our results suggest that the DMN constantly engages in online summarization and integration of paragraph-level incoming contextual information with information accumulated across minutes, hours, and even days. Such online summarization and integration provide the brain with the necessary capacity to flexibly integrate information accumulated over multiple timescales, a capacity that is currently lacking in the fixed contextual window architecture of many LLMs.

## Results

### Overview

The results and analyses are divided into three consecutive phases. First, we carried out a systematic analysis to investigate the effect of increasing the size of the incoming context window of the LLM’s input on the ability to predict the fMRI signals from the LLM’s embedding representations. For that, we applied the well-established neural-encoding analysis^7,8,10^ and tested its performances while varying the size of the context window from 8 tokens to the maximal possible size of 2048 tokens. In the second stage, we introduced our novel incremental context model, which combines both a short-term incoming context window and the long-term aggregated context. Then, we tested our model’s performance in predicting brain activity compared to a baseline LLM with either a long or short incoming context window. Lastly, in the third complementary stage, we performed a spectral analysis on the BOLD signal, for each brain area, to estimate how fast/slow the information changes – a measurement that is equivalent to estimating the amount of prior context the brain area processes in the present. Based on the results of this analysis, we identified brain areas that utilize long/short context windows and tested whether our incremental long-term context model predicts their activity better/worse than a short-term context model. All these stages are detailed in the following sections.

### Incoming Context is Processed in the Brain Through Small Context Windows

From the Narratives fMRI dataset^21^, we extracted data from 219 individuals who were passively listened to narrative stimuli. The data contained a total of 297 scans from 8 different relatively long stories/narratives (∼7 minutes or longer) which together encompass 15260 tokens (See Table S1 and Method). Brain images were all parcellated into 400 regions of interest (ROIs) according to the Schaefer’s atlas^22^ and then were averaged across participants in order to minimize as much potential noise as possible. Word embedding representations were created for each story using a state-of-the-art, open-source GPT3-like model (GPT-neoX^23^) and were subsequently used to predict the neural signals recorded from individuals who listened to that story, via the well-established neural-encoding analysis^10,7,24^ (see Method). We systematically tested the model’s predictions while varying the amount of prior context (i.e., the number of tokens) the model was exposed to during the word embedding extraction. We tested the following context window sizes: 8, 16, 32, 64, 128, 256, 512, and 2048 (the maximal possible value) tokens. The neural encoder model was trained and tested using 5-fold cross-validation for each window size and ROI separately, as detailed in the Method section.

The pattern of the results was clear. As Figure 2 presents, the performances of the neural encoder (as measured via the Pearson’s *r* correlation between the original and the predicted signal) are getting better as the window size increases, but only up to a window size of 32 tokens. From that point onwards, however, the performance tends to decrease as the window size increases, eventually reaching a plateau above 128 tokens. This pattern is reflected in terms of the number of ROIs where the *r*-score was significant (26, 129, 147, 87, 34, 34, 35, and 41 significant ROIs for 8, 16, 32, 64, 128, 256, 512, and 2048 tokens respectively), the maximal *r-*score values (0.13, 0.21, 0.29, 0.23, 0.16, 0.18, 0.18, and 0.19 for 8, 16, 32, 64, 128, 256, 512, and 2048 tokens respectively), and the averaged *r-*score across the 400 ROIs (0.04, 0.07, 0.08, 0.06, 0.03, 0.04, 0.04, and 0.04 for 8, 16, 32, 64, 128, 256, 512, and 2048 tokens respectively; See Figure 2b). These results are replicated almost identically when using the (relatively) older GPT-2 model, as presented in Figure S1. Moreover, in Figure S2 we show that the failure of large-context-window LLMs in predicting the brain is also observed with other LLMs that were designed specifically for long contexts: Long T5^25^, Transformer XL^26^, and Longformer^27^.

**Figure 2.**
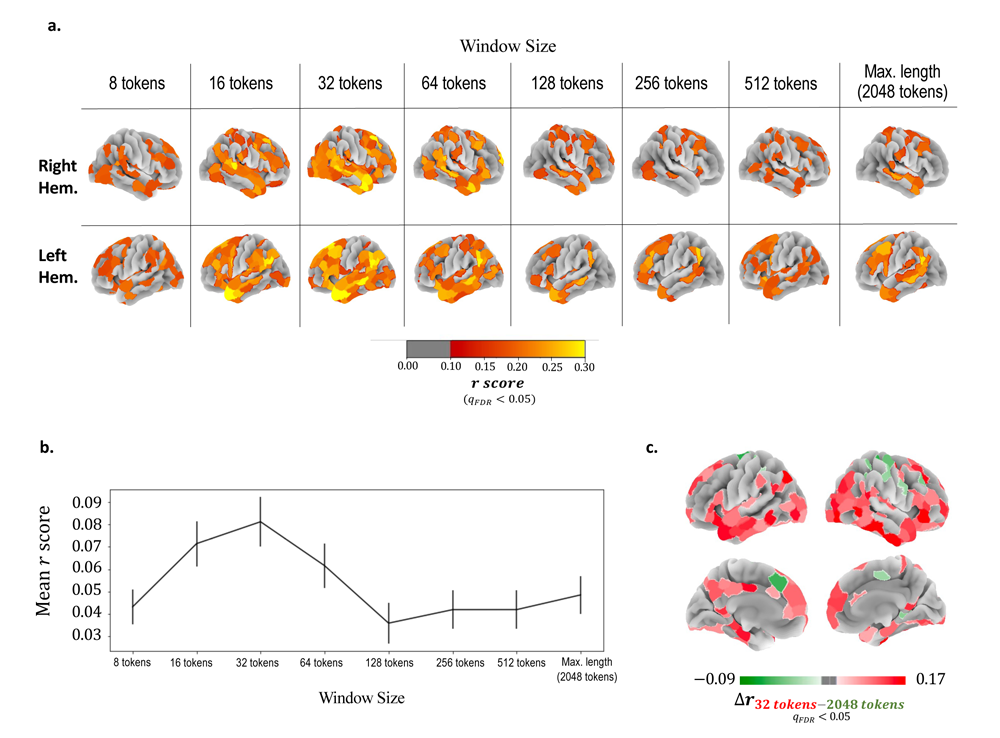
The effect of the size of the context window (in # of tokens) fed into the LLM on its ability to predict the neural signals. **a.** Cortical maps for different window sizes showing ROIs where the neural encoder score (Pearson’s *r* calculated between the predicted and the original signals) was statistically significant. **b.** The mean neural encoder *r*-score (averaged across ROIs) at different window sizes. Error bars represent the 95% confidence interval of the mean. **c.** A cortical map showing ROIs that were predicted significantly better using a window size of 32 tokens than with a window size of 2048 tokens (red areas), or vice versa (green areas). Note that in all of the green areas, although the *difference* between the *r*-scores was significantly in favor of the 2048-window size, the *r-*scores *themselves* were negligible and not significantly different from zero (none of the green areas are shown in figure 2a).

To further validate the results, we also conducted a direct comparison between the short window size of 32 tokens and the maximal long window size of 2048 tokens. For each ROI, we calculated *Δr_32 tokens-2048 tokens_* which equals to the *r*-score obtained from the LLM with a window size of 32 tokens, minus the *r*-score obtained from the LLM with the large window size of 2048 tokens. The vast majority of the significant ROIs in the resulting map were in favor of the short window size (103 ROIs with a significant positive *Δr_32 tokens-2048 tokens_* value; *Sig. threshold (Min)* = 0.054, *Max*= 0.17, *Mean* = 0.09, *SD* = 0.02; Figure 2c *red* areas). The resulting map also shows a few ROIs that were predicted significantly better by the long window size model than by the short window size model (15 ROIs with a significant negative *Δr_tokens-2048 tokens_* value; *Sig. threshold (Max)* = -0.04, *Min*= -0.09, *Mean* = -0.06, *SD* = 0.01; Figure 2c *green* areas). But it is important to note that although these areas achieved significant negative *Δr_tokens-2048 tokens_* values, the *r*-scores of the long window size model in these areas are lower than the statistical threshold (i.e., not significantly different from zero; note in Figure 2a that these areas are not presented).

The above results suggest that the success of fixed-size context LLMs in predicting language-related neural responses is effective only when the encoded information is related to a relatively short context window, equivalent to a timescale of several sentences. Moreover, as illustrated in Figure 2, this limitation was not only observed in temporal areas, which were previously linked to short timescales,^19,20^ but also identified in higher-order areas at the DMN associated with longer timescales. This confirms our first hypothesis that the incoming context to the brain is limited to small context windows containing up to tens of words, as the brain, unlike LLMs, cannot compute hundreds and thousands of tokens in parallel. In the next section, we present an alternative model that is capable of incorporating and maintaining very long contextual information in a sequential and incremental manner—similar to how we believe the human brain functions.

### An Alternative Cognitively Plausible Model for Short and Long Context Integration – The Incremental Context Model

The main limitation in using LLMs with large context windows to model long-term contextual computing in the human brain is the necessity to process hundreds of words in parallel. To cope with this limitation, we designed an alternative model where the window size of the input is kept (relatively) small, and yet contained both short-term and long-term contextual information. In this model, the context window comprises two components: incoming short-term context and aggregated long-term context. The incoming context consists of the last N words, where N is no more than several dozen tokens. The long-term component contains earlier information that appeared outside the narrower short-term window. Importantly, this information is no longer the original (hundreds or thousands) words from the stimulus, but instead a concise summary that is generated by the model itself.

Specifically, using a dedicated prompt design (see Method), we interact with the model and request it to generate a short summary based on previous information. This interaction was applied every several words, such that the summary is continuously updated and changed as the model advances through the story. Importantly, in order to capture long-term context, the summary of each step was generated based on the content of both the last several tokens, as well as the summary from the previous step (See figure 3b, figure S3b, and Method). In this way, the model always maintains an incremental long-term context through natural language text, which can then be used as input to the model again. As depicted in Figure 3b, a word embedding representation of a token is obtained from this model by concatenating two elements into its input: a short-term context window (consisting of the last 32 tokens; see the Method section for details on this choice) and the most recent state of the incremental summary, which provides the long-term context. A more detailed schematic illustration, including a textual example from the actual data, is presented in Figure S3.

**Figure 3.**
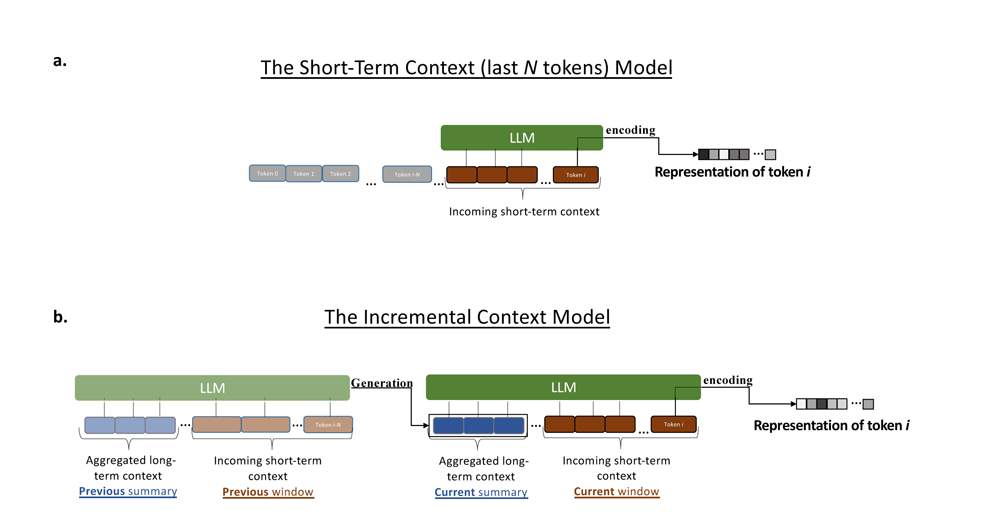
Schematic illustrations depicting the process of extracting contextual embedding representations for a single token using both the baseline short-term context model (a) and the novel incremental context model (b) **a.** The short-term, N-tokens (N=32) model. The word embedding representation of the token *i* is extracted by providing the LLM with that token, as well as the preceding N tokens. **b.** The long-term, incremental context model. To extract the word embedding representation for the same token *i*, the input to the model included both the short N-token window (as in the N-tokens model) and a concise natural-language summary generated by the model, which was based on information from the long-term context. This long-term contextual summary is updated as the model progresses through the story. It is generated based on the text that appeared before the short N-token window, as well as on the summary generated at the previous update step by the model. See Figure S3 for more details.

### The Incremental Context Model Better Predicts Neural Activity in Many Higher-Order Brain Areas

To assess the predictive power of the incremental model in modeling the integration between short-and long-term contexts in the human brain, we directly compared its performance in neural encoding against the following two baseline models: An LLM with a short-term incoming context window of 32 tokens (we chose 32 tokens as it was the optimal model in our first analysis. See Figure 2) and a “full-size” LLM with the maximum length of window size, i.e., 2048 tokens. Note that all the three models (i.e., Incremental-context, 32-tokens window size, and 2048-tokens window size) are based on the same pre-trained LLM and they differ only in the type/size of the context used in the input during the encoding phase (i.e., the extraction of the word-embedding representations).

Figure 4 presents the cortical maps of the comparisons between the models. For each pair of models, we calculated ΔΔ per ROI (out of 400 parcellations according to the Schaefer’s atlas^22^), which equals to the difference between the neural-encoder *r*-score of one model and of the other model. First, comparing to the 2048-tokens model, our novel incremental context model significantly improves the *r*-score in many parietal, temporal, and frontal ROIs, as reflected by positive *Δr_tokens-2048 tokens_* values (a total of 116 significant ROIs; *Sig. threshold (Min)* =0.05, *Max*= 0.17, *Mean* = 0.08, *SD*= 0.02; Figure 4a). Furthermore, none of the ROIs showed a significant negative *Δr_tokens-2048 tokens_* score, namely, there are no brain areas where their signal is predicted better by the 2048-tokens model comparing to the incremental context model. The results of this comparison demonstrate the substantial advantage of modeling long-term context in the brain using an incremental mechanism instead of parallel computing over hundreds of tokens.

**Figure 4.**
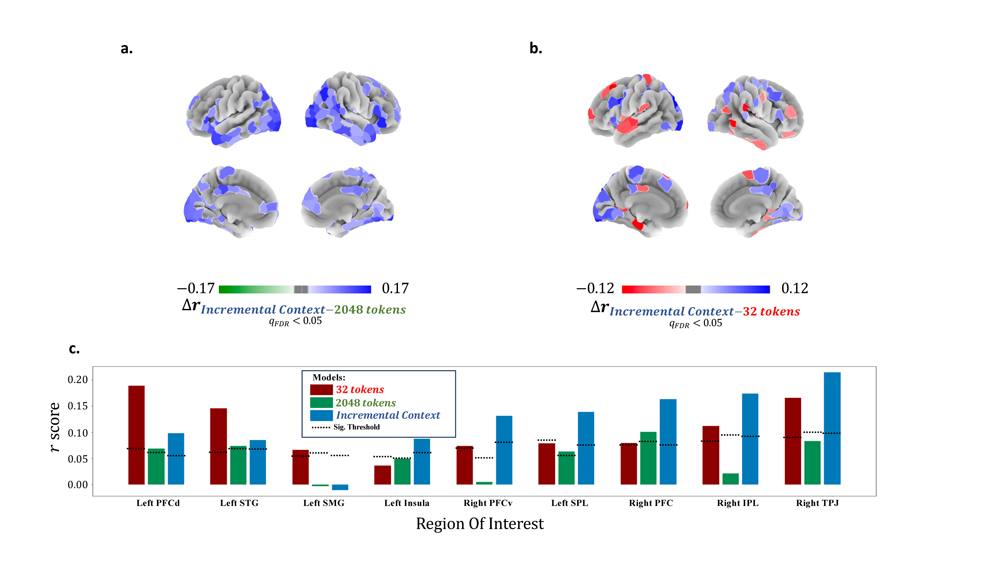
Comparisons between our incremental context model, the baseline long-term full-transformer (2048 tokens) model, and the baseline short-term context model (32 tokens). **a/b.** Cortical maps show the differences between the models in the neural encoder scores at different ROIs. The following comparisons were made: *Incremntal Context* vs. *2048 tokens* (a), and *Incremental Context* vs. *32 tokens* (b). The maps displaying only the ROIs that demonstrate a significant difference using a 10,000-iteration permutation analysis and FDR correction. **c.** Bar plots display the neural encoder results (y-axis) for selected language-related ROIs, depending on the model used for embedding representations (32-tokens/2048-tokens/Incremental Context). The horizontal dashed lines represent the statistical threshold resulting from the permutation analysis, i.e., only bars that are higher than this line are significantly different from zero. Notes. STG = Superior Temporal Gyrus; IPL = Inferior Parietal Lobule; SPL = Superior Parietal Lobule; TPJ = Temporoparietal Junction; SMG = Supramarginal Gyrus; PFC = Prefrontal Cortex; PFCd = Dorsal Prefrontal Cortex; PFCv = Ventral Prefrontal Cortex.

Second, we compared the results of our incremental context model to the results of the 32-tokens model, which according to our previous analysis (Figure 2) is the best model for the incoming *short-term* context. Figure 4b presents the cortical map of the *Δr_tokens-2048 tokens_* values and discovers both brain areas that are better predicted by the incremental model (positive *Δr_tokens-2048 tokens_* values), as well as brain areas that are better predicted by the short-term context model (negative *Δr_tokens-2048 tokens_* values). Our long-term context model (incremental context) significantly outperforms the short-term context model (32-tokens) in many areas among the Default Mode Network (DMN; 53 significant ROIs; *Sig. threshold (Min)* = 0.045, *Max*= 0.117, *Mean* = 0.06, *SD* = 0.01; Figure 4b *blue* areas). On the other hand, we also found brain areas that were predicted significantly better by the short-term context model (32-tokens), compared to the long-term incremental context model. This effect was mainly observed in the superior temporal gyrus, supramarginal gyrus, and areas in the dorsal prefrontal cortex (26 significant ROIs; *Sig. threshold (Max)* = 0.045, *Min*= -0.11, *Mean* = -0.06, *SD* = 0.01; Figure 4b *red* areas).

Importantly, the full cortical map of the *Δr_Incremental Context-32 tokens_* values (Figure 4b and Figure 4a) is aligned with the hierarchical structure of the time-scale changes discovered elsewhere,^19,20^ where the primary auditory cortex (located on the superior temporal gyrus) is involved in the most short-scale processing, and the more high-order DMN areas in the parietal lobule, as the TPJ and the precuneus, are associated with more long-term processing (Figure 1a). Moreover, the finding that DMN areas were better predicted by our incremental model, while the lower-level areas were better predicted by the short-incoming context LLM, is well aligned with our second hypothesis. That is, that higher-order DMN areas receive the incoming paragraph-level (i.e., short window of ∼32 tokens) contextual information from the downstream, lower-level areas, and engage in online summarization and integration of this information with information accumulated across the narrative (Figure 1b).

### A model-free spectral analysis of the frequency domain supports the model-based analyses

In this analysis we focused on the above presented cortical map of the contrast between the *r*-scores of our incremental model and the short-term context model (i.e., the *Δr_Incremental Context-32 tokens_* values; Figure 4b). As mentioned earlier, this map revealed a hierarchical organization of the cortex in terms of the timescale of the context processing and confirmed our hypothesis regarding the long-term contextual integration at the top level of the hierarchy. In Figure 5a we re-visualize this map, this time over the entire cortex (400 ROIs), with no statistical threshold (as in Figure 4b). This map demonstrates the gradient between the red areas – areas that are more associated with the short-term context (negative *Δr_Incremental Context-32 tokens_* values) – and the blue areas, which are more associated with the long-term context (positive *Δr_Incremental Context-32 tokens_* values). In the subsequent analysis, we conducted a complementary, model-free analysis on the fMRI (BOLD) signals, which provides additional validation for the timescale hierarchy revealed through the models.

**Figure 5.**
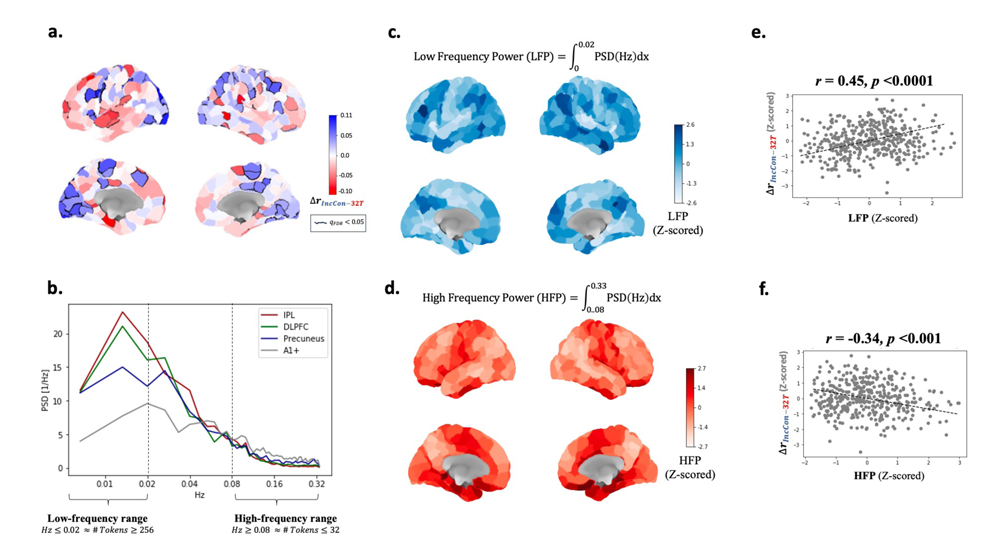
The complementary spectral analysis. **a.** The cortical map of the *Δr_Incremental Context-32 tokens_* (abbreviated as Δ*r_IncCon-32T_*) scores replicated from Figure 3b but with no statistical thresholding (significant ROIs are encircled here by a black line). This map shows the overall pattern of the cortical hierarchy between ROIs that are better predicted by the long-term Incremental-context model (more blue areas) and ROIs that are better predicted by the short-term (32 tokens) model (more red areas). **b.** The power spectral density (PSD) calculated for the BOLD signal within four important ROIs: Inferior Parietal Lobule (IPL), Dorsolateral Prefrontal Cortex (DLPFC), the Precuneus, and the primary auditory area. **c/d.** Cortical maps showing the power of high (c) and low (d) frequencies at different ROIs. These measurements were extracted from the PSD curve as described in the Method. **e/f.** Scatter plots representing the correlation between the *Δr_Incremental Context-32 tokens_* scores (figure 4a) and the high (e) or low (f) frequency power of the signal.

Timescale differences were investigated by looking at the frequency domain of the signal. Intuitively, when a signal mostly consists of low frequencies, it indicates that the encoded information changes slowly and gradually. As a result, the information at a certain time point does not deviate significantly from the information encoded far away in the past. Conversely, high frequencies in the signal suggest rapid changes of the information, and as such, the value of a certain time point is merely influenced by nearby time points, with less impact from distant past information. Therefore, performing a spectral analysis of the fMRI signal provides a reliable approximation of the extent of past contextual information processed by a certain brain area.^20^

For each ROI, we estimated the Power Spectral Density (PSD) of the averaged signal (See Method for detailed information), which illustrates how the power of the signal is distributed across different frequencies (see Figure 5b for selective ROIs). Next, we quantified the power of the high frequencies of the signal (which are equivalent to short context window sizes) by calculating the area under the PSD curve within the high frequencies range. Since the optimal size of the incoming context to the LLM is 32 tokens (Figure 2), we chose frequencies that demonstrate cycles (wavelengths) equivalent to the time taken to produce 32 tokens or less. According to our data, this time interval is approximately 12 seconds, and the corresponding frequency is ∼0.08 Hz (0.08*Hz* ≈ 1/12). Therefore, we calculated the integral of the PSD function between 0.08Hz and the Nyquist frequency, 0.33Hz, and denoted this value as the High Frequency Power (HFP) of the signal (*HFP* = ∫_0.08_^0.33^ *PSD*(*Hz*)*dx*; Figure 5c). Most importantly, we found a significant negative correlation between the HFP and the *Δr_Incremental Context-32 tokens_* values (*r* = -0.34, *p* < 0.001; Figure 5e). Namely, the higher the power of high frequencies within the ROI, the better it is predicted by the short-term context model compared to the long-term context model (i.e., a more ^negative ΔΔ^.(/0’1’($23 4%($’5$*!” $%&’() ^value).^

The above analysis was replicated for the low frequencies as well. We calculated the Low Frequency Power (LFP) of the ROI’s signal by taking the integral of the PSD between 0Hz to 0.02Hz (*LEP* = ∫_0_^0.02^ *PSD*(*Hz*)*dx*; Figure 5d). This range of frequencies is equivalent to window sizes of 256 tokens or longer (the cutoff of 256 tokens best demonstrates our results, but the overall pattern is preserved for different thresholds above 32 tokens as well). In contrast to the HFP, the LFP scores showed a strong *positive* correlation with the *Δr_Incremental Context-32 tokens_* values (*r* = 0.45, *p* < 0.0001; Figure 5f). This means that the greater the presence of low-frequency power in the ROI, the better it is predicted by the long-term incremental-context model compared to the short-term context model.

## Discussion

We hypothesized that, unlike the ability of current LLMs to process large contextual windows of hundreds and thousands of words in parallel, the human brain applies a different, more sequential, and flexible mechanism. In line with previous studies that demonstrated the topographical timescale hierarchy of temporal processing in the brain (Figure 1a)^18–20^, we proposed that (1) downstream, primary areas in the brain process the entire incoming context window up to tens of words (i.e., paragraph level), and that (2) higher-order areas in the DMN constantly engage in online summarization and integration of this short incoming contextual information with information accumulated across minutes, hours, and even days. First, we demonstrated that the default implementation of the incoming context window in LLMs, i.e., simply feeding the last *N* tokens into the model, can be considered a good model for the brain only when *N* is relatively small (*N*=32). However, when it comes to longer window sizes, the LLMs are no longer efficient in predicting the fMRI signal (Figure 2). This supports our first hypothesis that the brain can process the entire incoming context window only when the window contains no more than tens of words (equivalent to a paragraph-level).

Second, according to our second hypothesis, we proposed an alternative LLM-based incremental model for integrating long-term context information, beyond the recent tens of words. In contrast to feeding the entire text all at once to the model, our incremental context model preserves a modest number of tokens for parallel processing while retaining essential contextual information from tokens that were processed much earlier. By employing prompt-engineering techniques^28^, we have the model intermittently interact with the text and generate a natural language aggregated summarization that is integrated with the incoming short context window (Figure 3). Next, we empirically show that our novel incremental context model outperforms the alternative long-context-window LLM in predicting neural signals of long narratives (Figure 4). Moreover, in line with our second hypothesis, we found that among the DMN areas (located at the top level of the timescale hierarchy), the incremental model (which integrates both incoming and aggregated contexts) outperforms the short-(32 tokens) incoming-context window LLM. In contrast, the short-context LLM outperforms the incremental model in predicting lower-level brain areas located at the downstream of the timescale hierarchy (e.g., STG). Finally, we used a complementary spectral analysis to map cortical areas on a scale ranging from short-to long-term contextual processing by quantifying the power of low and high frequencies (the lower the frequency of the signal, the slower its fluctuations, and consequently, its timescale). Next, we demonstrated that the more dominant the brain area is in low frequencies, the better its signal was predicted by our long-term incremental context model. Similarly, when a brain area exhibited more high frequencies, it was predicted more accurately by the short-term context (32 tokens) implementation of the LLM (Figure 5).

Although the results of this study suggest that our incremental context model is a better fit for long-term context processing in the brain, it is not guaranteed that this model is completely cognitively plausible, but rather that it is more plausible than the default transformer model. First, it is not clear whether the long-term context is aggregated in the brain in the form of discrete words, as implemented in our model (i.e., through the generated summary), or rather in a more continuous way. Second, it is not clear whether aggregating information via summarization is, in fact, cognitively plausible. It could be that there are other, more cognitively plausible methods of aggregating contextual information rather than via summarization. As one way of justifying our proposed model as a cognitive mechanism, we have considered several alternative aggregation methods, such as extraction of key sentences from the earlier information or asking the model to generate leading keywords out of the text. All of these alternative aggregation methods were substantially inferior compared to our proposed summary generation model. Nevertheless, we do not rule out the possibility that future studies may uncover more effective alternatives.

This study is the first to use LLMs for modeling (very) long-term context processing in the human brain. Previous studies in the neural encoding literature all used LLMs that were only aware of the short-term context, i.e., where the context window size was as large as several tens of words.^1–10^ This was the case even though the participants in these studies were exposed to language stimuli that encompassed hundreds and even thousands of words. To the best of our knowledge, no study has investigated yet the effect of the LLM input contextual window size on the neural encoding performance. Moreover, no study has reported the neural encoder performance when the LLM is provided with a full-length context window. It is important to refer here to Caucheteux’s et al.’s^4^ work, where they empirically investigated the timescale hierarchy in the brain using LLMs. The current work is different for two reasons. First, unlike the current study, they did not manipulate the timescale by varying the number of tokens in the input; instead, they achieved this by scrambling the text at multiple levels (words, sentence etc.), similar to Lerner’s et al.’s work.^19^ Second, in all experimental conditions, they used a fixed window size of 256 tokens in the context. Consequently, the maximum timescale of processing they could investigate was limited to the size of one or two paragraphs. In our study, on the other hand, we primarily focused on very large timescales, spanning hundreds and thousands of tokens.

From the NLP-field perspective, our proposed incremental context model provides a new approach to dealing with the processing of very large texts.^29^ Apart from the cognitive plausibility gap addressed in this paper, the parallel computing nature of the transformer-based LLMs yields a quadratic computational complexity (*O*(*η*^“^)) which makes the processing of long texts significantly more expensive. Several solutions have been proposed in recent years to cope with this computational complexity problem. These included novel architectures of the attention matrix (e.g., sparse-attention^30^, dilated sliding window^27,29,31^, transient global attention^25^ and others), hierarchical combination of multiple transformers^9,32^, and implementations of RNN-like (recurrent neural network) modules within the transformer block.^26^ To the best of our knowledge, our incremental context model is the only model that instead of processing the entire text all at once, it repeatedly applies itself along the text, interacting with the information in a manner akin to human-like comprehension [note that the Transformer-XL model^26^ incorporates some form of recurrent processing within the transformer block, but it still takes the entire text as its input in parallel. In Figure S2, we empirically demonstrate that this model is indeed less effective at predicting human neural signals.]. While this study does not delve into assessing the incremental context model’s performance in various NLP tasks related to long texts, we hold the belief that its potential extends far beyond the field of cognitive neuroscience.

## Method

### FMRI data

The data for the present research was retrieved from the narratives dataset published elsewhere.^21^ The narratives dataset contains a variety of functional MRI datasets collected while human subjects listened to naturalistic spoken stories. Since the current work focuses on long-term context, we only gathered data of stimuli contain a single coherent long story (there are stories like “Schema” and “The 21^st^ Year” that contain multiple narratives within the story), and that do not contain any ambiguities or special experimental manipulations (such as “Shape” and “Green eyes”). Our final sample included data from 8 stories (“Lucy”, “Merlin”, “Pie man”, “Tunnel”, “Bronx”, “Sherlock”, “Not the fall”, and “Milky way”). It consists of 297 scans recorded from 219 individuals (78 individuals participated in more than a single stimulus), after excluding ’bad’ scans based on the publisher’s recommendation. Table 1 describes, for each story, the number of individuals, number of tokens in the story (according to the GPT’s tokenizer), and number of TRs. The publishers’ dataset paper reports the technical details regarding the MRI acquisition, as well as the entire preprocessing pipeline of the data.^21^ For our analyses, we downloaded the version of the data that was normalized to a surface template (the *fsaverage6* template of the FreeSurfer software^33^).

### Language model

All of the main analyses and models in this paper were based on the open-source, 20 billion parameters, GPT-NeoXT model (retrieved from https://huggingface.co/togethercomputer/GPT-NeoXT-Chat-Base-20B), developed by Together.ai and Eleuther.ai. This model is similar to Eleuther.ai’s GPT-NeoX model,^23^ but it has undergone further fine-tuning using a small amount of feedback data. According to the developers, GPT-NeoX and GPT-NeoXT are considered comparable to OpenAI’s GPT-3 and ChatGPT models, respectively. For replication and validation we also used the OpenAI’s GPT2 model^11^, as well as other LLMs designed for long-contexts, Long T5^25^, Transformer XL^26^, and Longformer.^27^

### Analyzing the effect of the context window size on neural encoding

*Story representations.* For each story in our dataset (out of 8 stories), we extracted word embedding representation for every single token. The word embedding representation is a 6144-dimensional vector provided in the last layer of the GPT-NeoXT model. To extract a representation for a token, the token is fed into the model together with tokens that appeared in the text prior to this token (i.e., the context window). We varied the number of tokens in the context window between 8, 16, 32, 64, 128, 256, 512, or 2048 (maximum length) tokens. This process yielded eight sets of word embedding representation vectors, corresponding to the eight window sizes. Each set takes the form of a *d* by *k* matrix, where *d* is the dimensionality of the word embedding vector and *k* is the number of tokens in the story, or the “time” dimension. To match to the time resolution of the fMRI data (which was sampled every TR = 1.5 seconds), we down-sampled the token-based time signal of the word embedding vectors to *n*, where *n* equals the number of TRs in the scan. This was done by averaging the vectors of all the tokens that appeared within each TR interval.

*Neural encoder model.* For each story, we first averaged the fMRI signal across participants in order to minimize the potential noise emerged from inherent individual differences. The BOLD signal was Z-normalized within voxel and participant prior to averaging the brains. Next, we parcellated the averaged cortical surface into 400 regions of interest (ROIs) using the parcellation atlas of Schaefer.^22^ Upon parcellation we again averaged the voxels’ signal within each ROI, yielding a total of 400 time course per story.

Subsequently, we constructed a neural encoder model, which is simply a linear regression model that predicts the neural signal, for each ROI, from the word embedding representation vectors. Formally, the neural encoder maps the embedding vectors matrix, *M*_7,(_, to the neural data matrix, *Y*_,++,(_, where *n* is the length of the story (i.e., number of TRs) and *d* is the dimensionality of the word embedding representation. Following previous studies^9,10^ we reduced the dimensionality of the word embedding representations, *d*, into 32 dimensions via principal component analysis (PCA). To train and test the neural encoder model, we applied 5-fold cross-validation. The story was split into five sections, and for each iteration, we left one section out for testing and trained the model on the remaining sections. The model was evaluated by calculating Pearson’s correlation coefficient (*r*) between the predicted signal and the actual brain signal of the test section. The *r*-scores obtained from the five folds were transformed to Fisher’s Z, averaged together, and then converted back to *r*-score. This process was separately done for each story. Since the number of participants was different in each story, we averaged the results from all the stories while assigning a different weight for each story as follows:

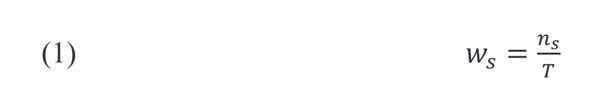

where *w*_*s*_ is the weight assigned for story *s*, *η*_*s*_ is the number of participants exposed to story *s*, and *T* is the total number of participants over the eight stories.

*Statistical testing.* To assess the significance of the results, we ran a permutation analysis with 10000 iterations. Foe each iteration, we randomly shuffled the phase of the neural signal and applied the above pipeline. The phase-shuffling method randomizes the signal while maintaining the mean and the autocorrelation of the original signal.^34^ It is implemented by applying the fast Fourier transformation on the original signal, randomizing only the phase component of the signal, and then applying an inverse fast Fourier transformation using the original frequency magnitudes and the randomized phases. The *r*-scores from all the 10000 permutations are saved and serve as the null distribution for the result. Based on this null distribution, we calculate the *p* value using the following formula:

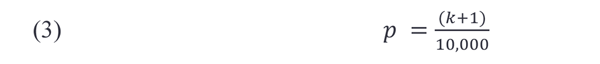

where *k* is the number of permutations that yielded a better result than the real result (i.e., the result of the unpermuted data). Correction for multiple hypothesis testing was applied using the false discovery rate (FDR) method.^35^

### The incremental context model

In the first analysis, we varied the context window size of the model and found that a window size of 32 tokens is optimal (Figure 2), and that increasing the context window is not beneficial in modeling long-term context. In our novel incremental context model the long-term context is not embodied with more tokens from the past, but rather with a compressed version of the past, which is a self-generated textual summary. Formally, when extracting the word embedding representation of a token *i*, the input to the model takes the following form:

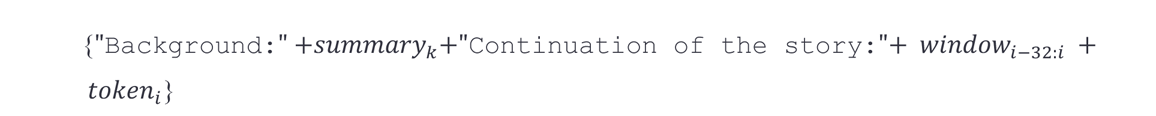

Specifically, the token *i* is concatenated with the preceding 32 tokens, denoted as *window_i-η’i_*, and before them, the *k*th (i.e., last updated, as described below) summary is presented. The strings “`Background:`” and “`Continuation of the story:`” are attached in the right places to help the model distinguish between the summary and the context window (see example in Figure S3). Note that *k* is different from *i* as the summary is not updated from toke to token, but for every 50 tokens. We adopted this slow updating method to (1) preserve a stable long-term summary and (2) because of the extreme computational and time resources that the model consumes during textual generation.

To obtain *summary_k_* (once in 50 tokens as described above), we used the pre-trained language-model head of the model to generate new text. The input to the model then was as follows:

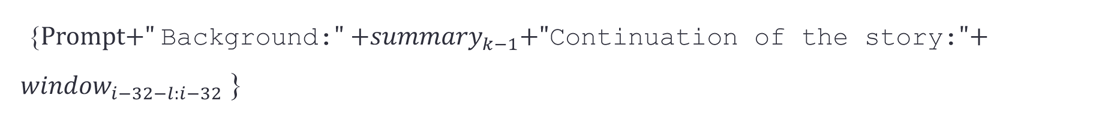

Where ’Prompt’ refers to our specifically designed textual instruction for the model, which was chosen after manual testing of several possibilities. The request prompt was “`Here are paragraphs taken from a story. Can you summarize the main theme in a few short sentences?`”. “*summary_k-1_*” represents the last summary generated at the previous step (we set *summary_0_* to an empty string). The expression “*window_i-32-l:i-32_*” refers to the text from the story that will be summarized along with the previous summary (Figure 3s). We tested multiple values of *l* (50, 100, 150, 200, and 250) by manually inspecting the generated text in some portions of the data (approximately 20%) and eventually set *l* = 100. Note from the subscript of “*window_i-32-l:i-32_*” that the text for summarization does not include the last 32 tokens, as this window is preserved intact in the input to the model when extracting the word embedding of token *i*, as described above.

In the summary generation we limited the length of the generated text to a maximum of 50 tokens. As a result, the total length of the input to the model during word embedding extraction was never greater than 90 tokens (the last token + 32 tokens of *window*_i-32:j_ + 50 tokens of *summary_k_* + 7 tokens of the strings in between). Importantly, this method allows us to provide the model with long-term context while preserving a reasonable number of tokens to be computed in parallel. We used the recommended method for text generation in this model^23^ which was multinomial sampling with temperature of 0.9.

### Direct comparisons between the models

To test the performance of the Incremental context model in neural encoding, we directly compare its *r* scores to two baseline models. One baseline was a short-term context model that takes only the last 32 tokens as its input (i.e., *window_j-32:i_* + *token_j_η*_’_}) with no additional long-term information. The other baseline model was a long-term context model that does not compress the long-term information but takes in its input all the tokens preceding the current tokens (up to the maximal possibility of 2048 tokens; i.e.,{*window_j-2048:j_* + *token_j_η*}).

For each pair of models {Incremental context vs. 32 tokens, Incremental context vs. 2048 tokens, *32 tokens* vs. *2048 tokens*}, we subtracted the *r* scores of one model (400 *r*-scores in total, corresponding to the 400 ROIs) from the *r*-scores of the other model and denoted the results as ΔΔ_!“#$% ’(!“#$% *s*_. In total, we calculated the following ^scores: ΔΔ^_*+,-$!$+.’%/“+.$0.(12.“3$+4_^, ΔΔ^_*+,-$!$+.’%/”+.$0.(2567 .“3$+4_^, and^ ΔΔ_12.“3$+4(2567.“3$+4_. To test the statistical significance of the results, we applied random phase-shifting permutation analysis, as described above (in the section titled “*Analyzing the effect of the context window size on neural encoding*”), calculated *p* values using formula (2) and applied the FDR correction.

### Correlating low frequency and high frequency power with

*Δr_Incremental Context-32 tokens_*

The averaged parcellated surface data (i.e. 400 ROIs) obtained from participants in each story where first analyzed through a spectral analysis. After Z-normalizing the signal to mean = 0 and variance = 1, we calculated the power spectral density (PSD) of the signal using the Welch method^36^ with a Hann window of 100 seconds and 50% overlap. The PSD curve describes how the power in a signal is distributed across different frequencies in the spectrum. Specifically, it is well-suited to our data as it enables the processing of signals with varying durations. Given that we have eight different stories, each with its unique duration, this method allows us to normalize the power to units of Hz in a consistent manner.

From the PSD curve, we derived the low frequency power (LFP) and the high frequency power (HFP) scores as follows:

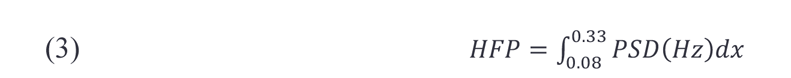

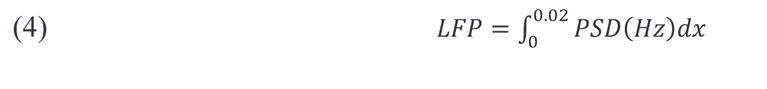

The lower bound for the integral of HFP is 0.08Hz because it is equivalent to a wavelength (cycle) of 32 tokens (32 tokens takes ∼12s, and therefore, 0.08Hz ≈ 1/12), and the higher bound, 0.33Hz, is the Nyquist frequency (the maximum decomposed frequency). The range of the integral on LFP starts form Zero (the minimum decomposed frequency) to 0.02Hz, which is equivalent to a window of 256 tokens. 0.02Hz was chosen because it maximizes the correlation between LFP and Δ*r_Incremental Context-32 tokens_*, however, the pattern of the results is preserved for different thresholds as well.

HFP and LFP scores were subsequently weighted averaged across the stories (as we did in the section “*Analyzing the effect of the context window size on neural encoding*”) using formula (1) above. Last, we calculated the correlations between Δ*r_Incremental Context-32 tokens_* and both the HFP and the LFP scores, over the 400 ROIs, using Pearson’s *r*.

## Supplementary materials

**Table S1.** Descriptive statistics of the data used in this study.

| Narrative | # of participants | # of tokens | # of TRs |
| --- | --- | --- | --- |
| **Lucy** | 14 | 1585 | 351 |
| **Merlin** | 35 | 2134 | 612 |
| **Pie man** | 73 | 954 | 286 |
| **Tunnel** | 21 | 3309 | 1013 |
| **Bronx** | 47 | 1356 | 374 |
| **Sherlock** | 36 | 2509 | 723 |
| **Not the fall** | 54 | 1561 | 383 |
| **Milky way** | 17 | 1052 | 282 |
| **Total** | 219 individuals | 15260 | 4314 |

**Figure S1.**
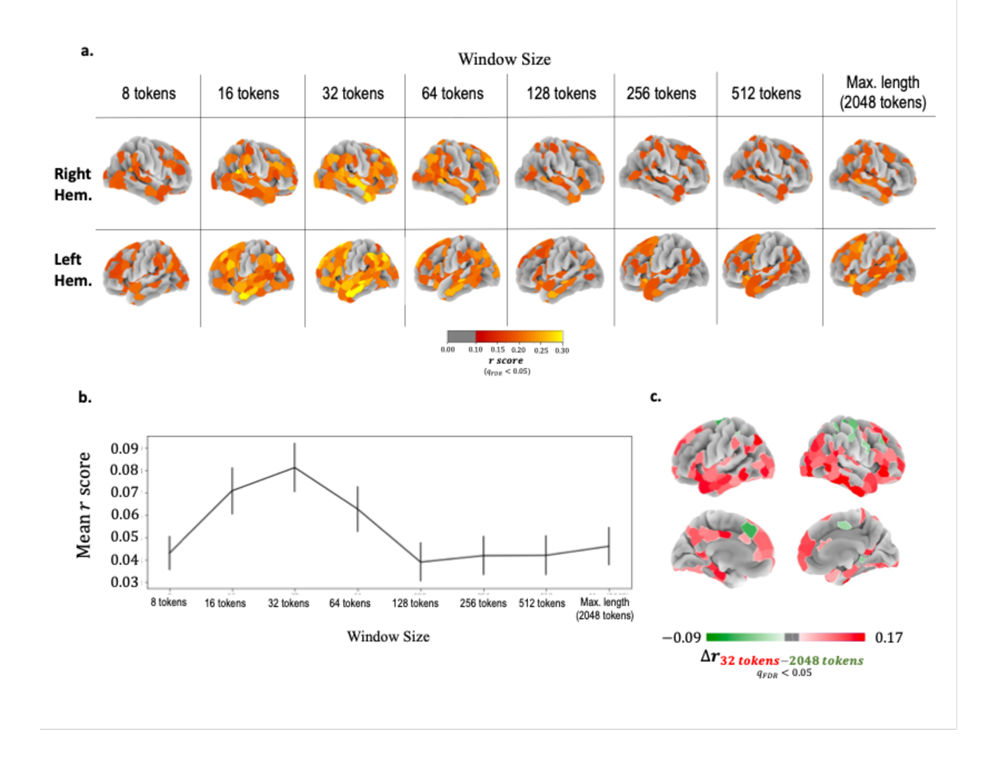
Replication of Figure 2 using the GPT-2 model.

**Figure S2.**
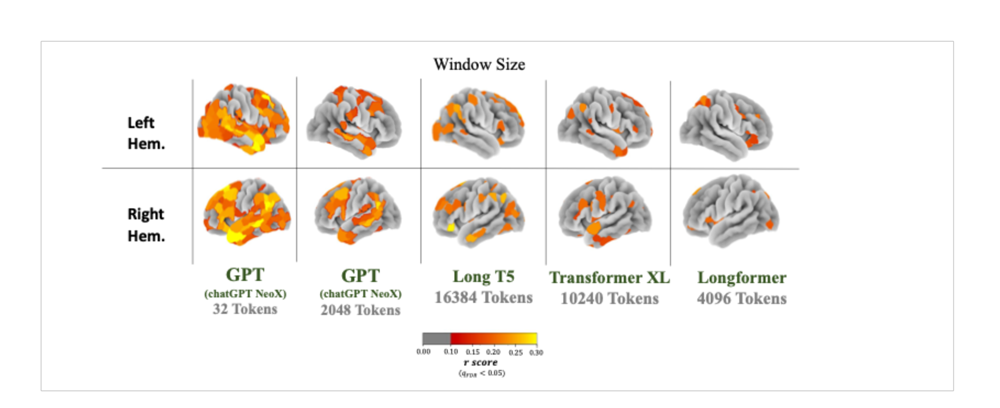
Cortical maps for different models/window sizes showing ROIs where the neural encoder score (Pearson’s *r* calculated between the predicted and the original signals) was statistically significant. The pattern shown in Figure 2a is replicated here. When the window size of the transformer model exceeds 32 tokens, the neural encoder’s performance decreases. This holds true even for models specifically designed for processing long texts.

**Figure S3.**
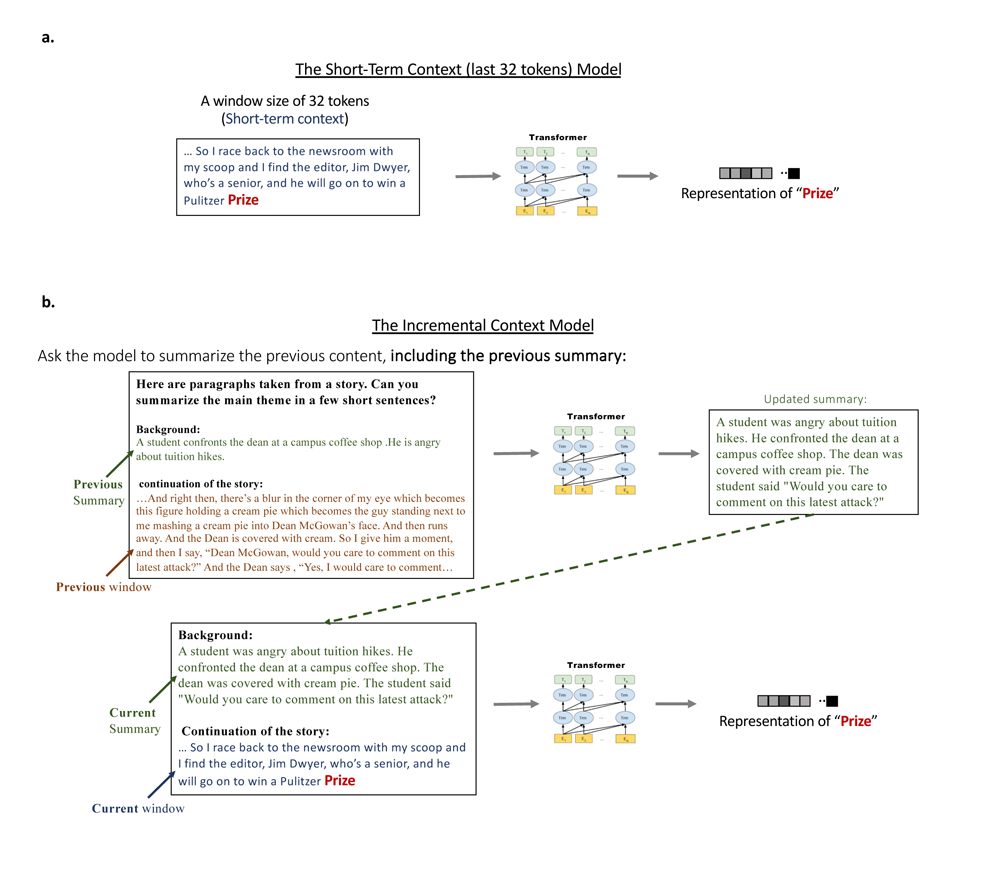
Schematic illustrations depict the process of extracting contextual embedding representations for a single token using both the baseline short-term context model (a) and the novel incremental context model (b) **a.** The short-term 32-tokens model. The word embedding representation of the token “*Prize*” is extracted by providing the LLM with that token, as well as the preceding 32 tokens. **b.** The long-term incremental context model. To extract the word embedding representation for the same token (“Prize”), the input to the model included both the short 32-token window (as in the 32-tokens model) and a concise natural-language summary generated by the model, which was based on information from the long-term context. This long-term contextual summary is updated as the model progresses through the story. It is generated based on the text that appeared before the short 32-token window, as well as on the summary generated at the previous update step by the model.

